# Nonsense mediated decay and a novel protein Period-2 regulate *casein kinase I* in an opposing manner to control circadian period in *Neurospora crassa*

**DOI:** 10.1101/2020.10.29.360735

**Authors:** Christina M. Kelliher, Randy Lambreghts, Qijun Xiang, Christopher L. Baker, Jennifer J. Loros, Jay C. Dunlap

## Abstract

Circadian clocks in fungi and animals are driven by a functionally conserved transcription-translation feedback loop. In *Neurospora crassa*, negative feedback is executed by a complex of Frequency (FRQ), FRQ-interacting RNA helicase (FRH), and Casein Kinase I (CKI), which inhibits the activity of the clock’s positive arm, the White Collar Complex (WCC). Here, we show that the *period-2* gene, whose mutation is characterized by recessive inheritance of a long 26-hour period phenotype, encodes an RNA-binding protein that stabilizes the *ck-1a* transcript, resulting in CKI protein levels sufficient for normal rhythmicity. Moreover, by examining the molecular basis for the short circadian period of *period-6* mutants, we uncovered a strong influence of the Nonsense Mediated Decay pathway on CKI levels. The finding that circadian period defects in two classically-derived *Neurospora* clock mutants each arise from disruption of *ck-1a* regulation is consistent with circadian period being exquisitely sensitive to levels of *casein kinase I.*

## Introduction

The *Neurospora* circadian oscillator is a transcription-translation feedback loop that is positively regulated by the White Collar Complex (WCC) transcription factors, which drive expression of the negative arm component Frequency (FRQ). The circadian negative arm complex is composed of FRQ, FRQ-Interacting RNA helicase (FRH), and Casein Kinase I (CKI), which promote WCC phosphorylation on key phospho-sites to inhibit its activity (Wang et al., 2019). Although not related by sequence similarity, FRQ is functionally homologous to PERs and CRYs in the animal clock. FRQ, PERs, and CRYs are extensively regulated transcriptionally, translationally, and post-translationally over the circadian day, and these regulatory mechanisms are highly conserved among clocks in animals and fungi (reviewed in: (Hurley et al., 2016)).

Negative arm components are regulated at the RNA and protein levels to maintain circadian phase and period. Anti-sense transcription at the *frq* locus produces the *qrf* transcript, which is required for proper phase control and light responses of the fungal clock (Kramer et al., 2003). The mammalian PER2 anti-sense transcript displays nearly identical dynamics to *qrf* expression (Koike et al., 2012). PER2 sense expression levels are further regulated by microRNA binding sites in its 3’ UTR (Yoo et al., 2017). In a similar manner, *frq* RNA is directly targeted for turnover by rhythmic exosome activity in the late day (Guo et al., 2009). Splicing of the *frq* transcript is regulated by temperature (Colot et al., 2005), mirroring thermal regulation mechanisms in the clocks of *Drosophila* (Majercak et al., 1999) and *Arabidopsis* (James et al., 2012). FRQ is an intrinsically disordered protein encoded by non-optimal codons to improve its co-translational folding (Zhou et al., 2013), and FRQ structure is also stabilized by its binding partner FRH (Hurley et al., 2013). PER2 is also largely intrinsically disordered, and indeed circadian clock proteins across species have large stretches of intrinsic disorder which are in the early stages of functional characterization (Pelham et al., 2020; Pelham et al., 2018) (reviewed in: (Partch, 2020)). Finally, an extremely conserved feature of the clock’s negative arm is progressive phosphorylation and alteration of function over time (reviewed in: (Dunlap and Loros, 2018)). FRQ is progressively phosphorylated over the day (Baker et al., 2009), as are CRY1 and PER2 in the mammalian oscillator (Ode et al., 2017; Vanselow et al., 2006). A conserved phospho-switch in mammalian PER2 and fly PER proteins has been implicated in both temperature compensation and in closing the circadian feedback loop (Philpott et al., 2020; Top et al., 2018). Taken together, FRQ, PERs, and CRYs are tightly regulated and underlying mechanisms are often conserved between clock models.

In contrast, less is known about the mechanisms regulating expression of the other essential member of the negative arm complex, CKI, orthologs of which are highly conserved in sequence and in function across eukaryotic clocks. CKI forms a stable complex as FRQ-FRH-CKIa in *Neurospora* (Baker et al., 2009; Gorl et al., 2001), as PER-DOUBLETIME (DBT) in flies (Kloss et al., 2001), and as a multi-protein complex of PER-CRY-CKIδ in mouse (Aryal et al., 2017). Fungal CKI phosphorylates both FRQ and WCC (He et al., 2006). Insect DBT and mammalian CKIδ/ε are key regulators of the PER2 phospho-switch, including phosphorylation of hPER2 at S662, which is associated with the human sleep and circadian disorder FASPS (Narasimamurthy et al., 2018; Philpott et al., 2020; Toh et al., 2001; Zhou et al., 2015). Significantly, mutation of human CKIδ itself phenocopies this, also leading to FASPS (Xu et al., 2005). CKI phosphorylations contribute to feedback loop closure in all species, and FRQ-CKI binding strength is a key regulator of period length in *Neurospora* (Liu et al., 2019). CKI abundance is not rhythmic in any species described to date (Gorl et al., 2001; Kloss et al., 2001), but preliminary evidence suggests that its expression levels are tightly controlled to keep the clock on time, just like FRQ/PER/CRY. In mammals, CKI knockdown or knock out significantly lengthens period (Isojima et al., 2009; Lee et al., 2009; Tsuchiya et al., 2016), and CKIδ levels are negatively regulated by m6A methylation (Fustin et al., 2018). In *Neurospora*, decreasing the amounts of the *casein kinase I (ck-1a)* transcript using a regulatable promoter leads to long period defects up to ~30 hours (Mehra et al., 2009). CKI has a conserved C-terminal domain involved in autophosphorylation and inhibition of kinase activity (Guo et al., 2019). Fungal mutants lacking this CKI C-terminal inhibitory domain have hyperactive kinase activity (Querfurth et al., 2007) and display short periods. Across clock models, the circadian period is sensitive to CKI abundance and activity due to its importance in circadian feedback loop closure.

Our modern understanding of the circadian clock was founded on genetic screens and characterization of mutants with circadian defects (Feldman and Hoyle, 1973; Konopka and Benzer, 1971; Ralph and Menaker, 1988). The fungal clock model *Neurospora crassa* has been a top producer of relevant circadian mutants due to its genetic tractability, ease of circadian readout, and functional conservation with the animal circadian clock (reviewed in: (Loros, 2020)). Forward genetic screens used the *ras-1^bd^* mutant background (which forms distinct bands of conidiophores once per subjective night) in race tube assays to identify key players in the circadian clock (Belden et al., 2007; Feldman and Hoyle, 1973; Sargent et al., 1966). Genetic epistasis among the *period* genes, and in some cases, genetic mapping of mutations was also performed using *N. crassa* (Feldman and Hoyle, 1976; Gardner and Feldman, 1981; Morgan and Feldman, 2001).

All but one of the extant *period* genes in *Neurospora* have been cloned, and their identities have expanded our knowledge of core-clock modifying processes. *period-4*, or Checkpoint Kinase 2 (Chk2), linked the clock to cell-cycle progression (Pregueiro et al., 2006). *period-3*, or Casein Kinase II (CKII), implicated direct phosphorylation of core clock proteins as central to temperature compensation (Mehra et al., 2009). *period-1* is an essential RNA-helicase that regulates the core clock under high nutrient environments (Emerson et al., 2015). *period-6* is a core subunit of the Nonsense-Mediated Decay (NMD) complex (Compton, 2003), although its circadian role remains cryptic. Among the available *period* genes, only *period-2* remains uncharacterized.

We have mapped the *period-2* mutation to NCU01019 using whole genome sequencing, and discovered its molecular identity; however, attributing its long period mutant phenotype to molecular function has remained elusive (Lambreghts, 2012). Equipped with the identity of PRD-2, we then followed up on the observation that the *prd-6* short period phenotype is completely epistatic to the *prd-2* mutant’s long period (Morgan and Feldman, 1997, 2001). We find that PRD-6 and PRD-2 use distinct mechanisms to play opposing roles in regulating levels of the *casein kinase I* transcript in *Neurospora*, thus rationalizing the circadian actions of the two clock mutants whose roles in the clock were unknown. PRD-2 stabilizes the *ck-1a* mRNA transcript, and the clock-relevant domains and biochemical evaluation of the PRD-2 protein indicate that it acts as an RNA-binding protein. We genetically rescue the long period phenotype of *prd-2* mutants by expressing a hyperactive CKI allele and by titrating up *ck-1a* mRNA levels using a regulatable promoter. The endogenous *ck-1a* transcript has a strikingly long 3’ UTR, indicating that its mRNA could be subject to NMD during a normal circadian day. We confirm that *prd-6* mutants have elevated levels of *ck-1a* in the absence of NMD, and further rescue the short period defect of *prd-6* mutants by titrating down *ck-1a* mRNA levels using an inducible promoter. Taken together, a unifying model emerges to explain the action of diverse *period* mutants, where the *casein kinase I* transcript is subject to complex regulation by NMD and an RNA-binding protein, PRD-2, to control its gene expression and maintain a normal circadian period.

## Results

### An Interstitial Inversion Identifies *prd-2*

Genetic mapping and preliminary analyses identified *period-2* as a recessive mutant with an abnormally long ~ 26 hour period length that mapped to the right arm of LG V (Morgan and Feldman, 1997, 2001). Genetic fine structure mapping using selectable markers flanking *prd-2*, in preparation for an anticipated chromosome walk, revealed an extensive region of suppressed recombination in the region of the gene, consistent with the existence of a chromosome inversion (Lambreghts, 2012). PCR data consistent with this prompted whole genome sequencing that revealed a 322 kb inversion on chromosome V (Lambreghts, 2012) in the original isolate strain hereafter referred to as *prd-2^INV^.* The left breakpoint of the inversion occurs in the 5’ UTR of NCU03775, and its upstream regulatory sequences are displaced in the *prd-2^INV^* mutant. However, a knockout of NCU03775 (FGSC12475) has a wild-type circadian period length, unlike the long period *prd-2^INV^* mutant (Supplementary Figure 1). The next closest gene upstream of the left inversion is NCU03771, but its transcription start site (TSS) is > 7 kb away. The right breakpoint of the inversion occurs in the 5’ UTR of NCU01019, disrupting 333 bases of its 5’ UTR and its entire promoter region (Figure 1A – B). A knockout of NCU01019 has a 26 hour long period, matching the *prd-2^INV^* long period phenotype (Figure 1C). The *prd-2^INV^* mutant has drastically reduced levels of NCU01019 gene expression in constant light and in the subjective evening (Figure 1D), suggesting that the inversion completely disrupts the NCU01019 promoter and TSS. Placing NCU01019 under the nutrient-responsive *qa-2* promoter, we find that the long period length occurs at very low gene expression levels using 10^−6^ M quinic acid induction (Figure 1E). Finally, ectopic expression of NCU01019 at the *csr-1* locus in the *prd-2^INV^* background rescues the long period phenotype (Figure 1F). We conclude that PRD-2 is encoded by NCU01019.

**Figure 1.**
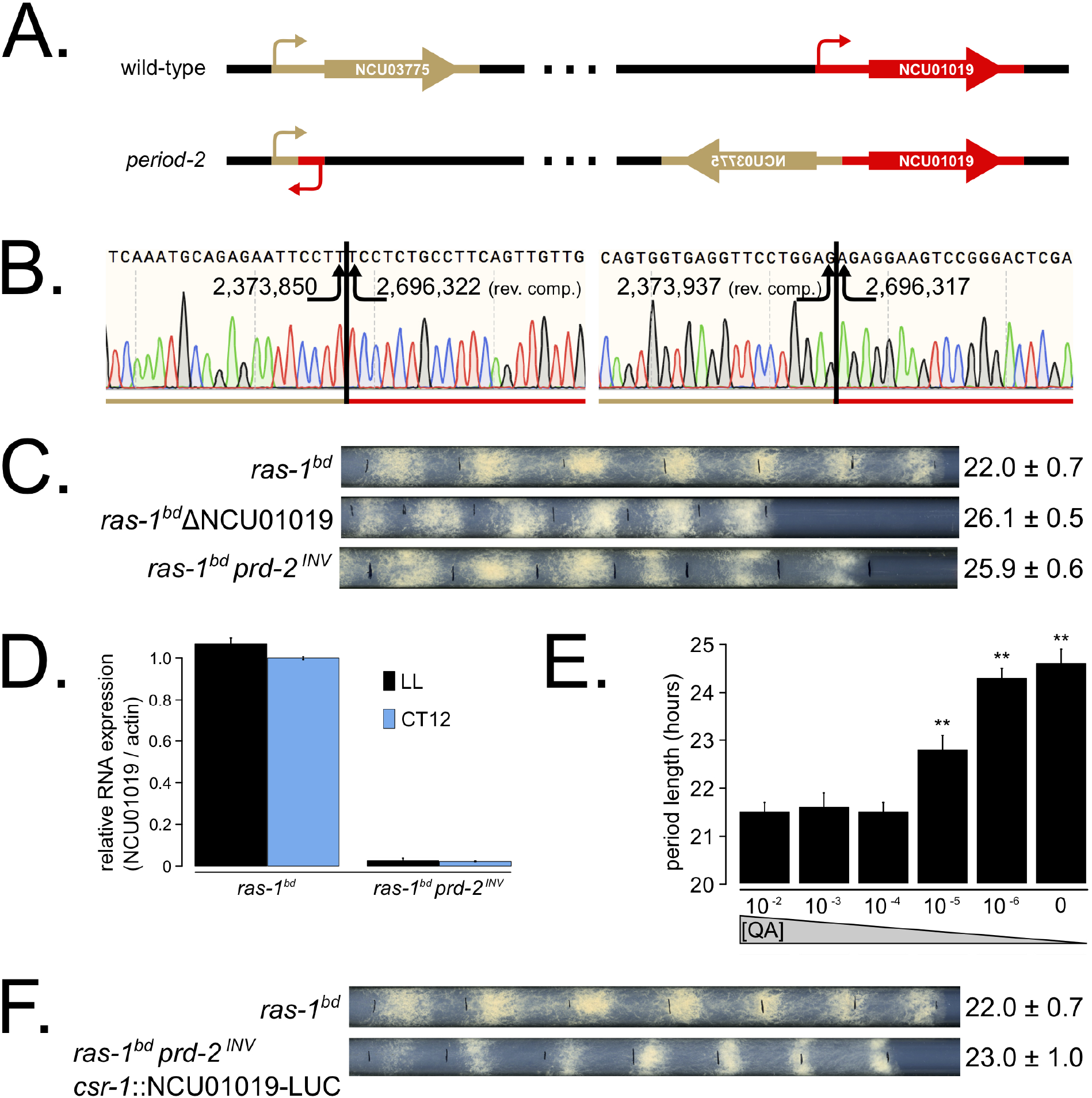
The *period-2* phenotype derives from reduced expression of NCU01019. Whole genome sequencing identified a 322,386 bp inversion on linkage group V in the original *period-2* mutant strain (Lambreghts, 2012). The inversion breakpoints disrupt two loci, NCU03775 and NCU01019, depicted in cartoon form (**A**). Sanger sequencing confirms the DNA sequence of the left and right breakpoints, and the corresponding NC12 genome coordinates are shown at each arrowhead (**B**). Circadian period length was determined by race tube assay for *ras-1^bd^* controls, targeted deletion of the NCU01019 locus, and the classically derived *prd-2^INV^* mutant. The ΔNCU01019 mutant has a long period and slow growth defect similar to *prd-2^INV^* (**C**). NCU01019 RNA expression levels are detectable by RT-qPCR in the *prd-2^INV^* mutant but are drastically reduced compared to *ras-1^bd^* controls grown in constant light or at subjective dusk CT12 (**D**). After replacing the endogenous promoter of NCU01019 with the inducible *qa-2* promoter, addition of high levels of quinic acid (10^−2^ – 10^−3^ M) led to a normal circadian period by race tube assay (10^−2^ M τ = 21.5 ± 0.2 hours; 10^−3^ M τ = 21.6 ± 0.3 hours; 10^−4^ M τ = 21.5 ± 0.2 hours). Lower levels of QA inducer led to a long circadian period (10^−5^ M τ = 22.8 ± 0.3 hours; 10^−6^ M τ = 24.3 ± 0.2 hours; 0 QA τ = 24.6 ± 0.3 hours) due to reduced NCU01019 expression. Asterisks (**) indicate p < 1×10^−10^ by student’s t-test compared to 10^−2^ M QA race tube results (**E**). The entire NCU01019 locus (plus 951 bases of its upstream promoter sequence) was fused in-frame with codon-optimized luciferase. Ectopic expression of this NCU01019-luc construct in the *prd-2^INV^* background rescues the long period phenotype by race tube assay (**F**).

We mapped the clock-relevant domains of the PRD-2 protein (Figure 2A), finding that both an SUZ domain and the proline-rich C-terminus of PRD-2 are required for a normal clock period. This result was confirmed in two separate genetic backgrounds either by replacing the endogenous locus with domain deletion mutants (Figure 2B) or by ectopic expression of domain mutants at the *csr-1* locus in a *Δprd-2* background (Figure 2C) (Supplementary Table 1). The SUZ domain family can bind RNA directly *in vitro* (Song et al., 2008), but curiously PRD-2’s adjacent R3H domain, which is better characterized in the literature as a conserved RNA-binding domain, is dispensable for clock function. The C-terminus of PRD-2 is predicted to be highly disordered, and finer mapping of this region showed that neither a glutamine/proline-rich domain (amino acids 525 – 612, 21% Gln, 26% Pro) nor a domain conserved across fungal orthologs (amino acids 625 – 682, 21% Pro) were required for normal clock function (Figure 2C). The remainder of the C-terminus (amino acids 495 – 524, 28% Pro; 683 – 790, 24% Pro) contains a clock relevant region of PRD-2 based on deletion analyses. Further, PRD-2 SUZ domain and C-terminal deletion mutants are expressed at the protein level, indicating that clock defects must be due to the absent domain (Supplementary Figure 2A). PRD-2 is exclusively localized to the cytoplasm based on biochemical evaluation, and this localization does not change as a function of time of day (Figure 2D).

**Figure 2.**
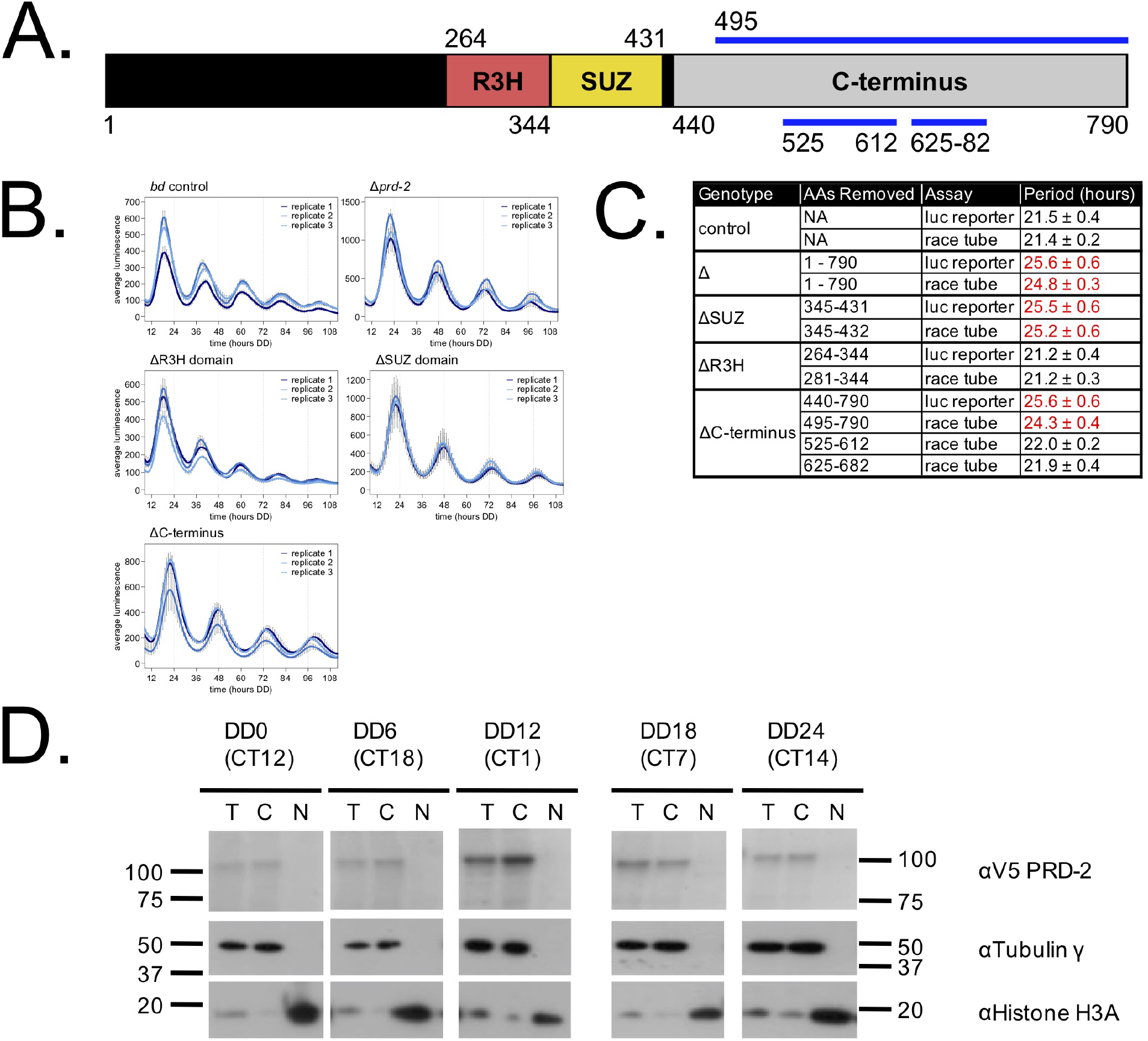
Clock-relevant protein domains and localization of PRD-2 suggest and RNA-binding function. PRD-2 has tandemly arrayed R3H and SUZ domains associated with RNA binding proteins, and its C-terminal region is highly enriched for proline (P) and glutamine (Q). The cartoon of PRD-2 protein lists relevant amino acid coordinates (**A**). The native NCU01019 locus was replaced with single domain deletion mutants, and 96-well plate luciferase assays were used to measure the circadian period length in triplicate wells per biological replicate experiment. A wild-type clock period was recovered in *ras-1^bd^* controls and the *prd-2ΔR3H* mutant, while *Δprd-2, prd-2ΔSUZ*, and *prd-2*ΔC-terminus had long period phenotypes (**B**). Independently constructed strains targeted domain deletion mutants to the *csr-1* locus in a *Δprd-2* background (Supplementary Table 1), and mutant period lengths were determined by race tube assay. Period lengths (± 1 SD) show that the clock-relevant domains of PRD-2 are the SUZ domain and the C-terminus (**C**). Total (T), Nuclear (N), and Cytosolic (C) fractions were prepared over a circadian time course (N = 1 per time point). g-Tubulin (NCU03954) was used as a control for cytoplasmic localization and histone H3 (NCU01635) for nuclear localization. PRD-2 tagged with a C-terminal V5 epitope tag is localized to the cytoplasm throughout the circadian cycle (**D**).

NCU01019 RNA expression is not induced by light (Wu et al., 2014) nor rhythmically expressed over circadian time (Hurley et al., 2014). NCU01019 protein is abundant and shows weak rhythms (Hurley et al., 2018) (Supplementary Figure 2B), which suggests that PRD-2 oscillations are driven post-transcriptionally to peak in the early subjective morning, prior to the peak in the *frq* transcript (Aronson et al., 1994). Rhythms in PRD-2 protein expression were confirmed using a luciferase translational fusion (Supplementary Figure 2C), which peaked during the circadian day. *prd-2^INV^* and ΔNCU01019 have a slight growth defect (Figure 1C) and are less fertile than wild-type as the female partner in a sexual cross (data not shown). Temperature and nutritional compensation of ΔNCU01019 alone are normal (Supplementary Figure 3), which was expected given the normal TC profile of the *prd-2^INV^* mutant (Gardner and Feldman, 1981).

### PRD-2 Regulates CKI Levels

To identify the putative mRNA targets of PRD-2, we performed total RNA-Sequencing on triplicate samples of *Δprd-2* versus control grown in constant light at 25°C. Hundreds of genes are affected by loss of PRD-2, but we did not identify a consensus functional category or sequence motif(s) for the putative PRD-2 regulon (Supplementary Figure 4). Given the pleotropic phenotypes of *Δprd-2*, we posit that PRD-2 plays multiple roles in the cell, including regulation of carbohydrate and secondary metabolism. Focusing specifically on core clock genes, we found that *ck-1a, frq, wc-2, ckb-1* (regulatory beta subunit of CKII), and *frh* were significantly altered in the absence of PRD-2 (Figure 3A). Pursuing the top two hits, we found that the CKI transcript was dramatically less stable in *Δprd-2* (Figure 3B), while *frq* mRNA stability was not significantly altered (Supplementary Figure 5). To demonstrate that PRD-2 binds the *ck-1a* transcript *in vivo*, we used RNA immunoprecipitation after UV crosslinking (CLIP). The Pumilio family RNA-binding protein PUF4 (NCU16560) was previously shown to bind in the 3’ UTR of *cbp3* (NCU00057), *mrp-1* (NCU07386), and other target genes identified by HITS-CLIP high-throughput sequencing (Wilinski et al., 2017). C-terminally tagged alleles of PRD-2, PUF4, and an untagged negative control strain were used to immunoprecipitate crosslinked RNAs (Materials & Methods). As expected, *cbp3* and *mrp-1* positive controls were significantly enriched in the PUF4 CLIP sample compared to the negative IP (Figure 3C). *ck-1a* is also enriched in the PRD-2 CLIP sample, demonstrating that the CKI transcript is a direct target of the PRD-2 protein (Figure 3C).

**Figure 3.**
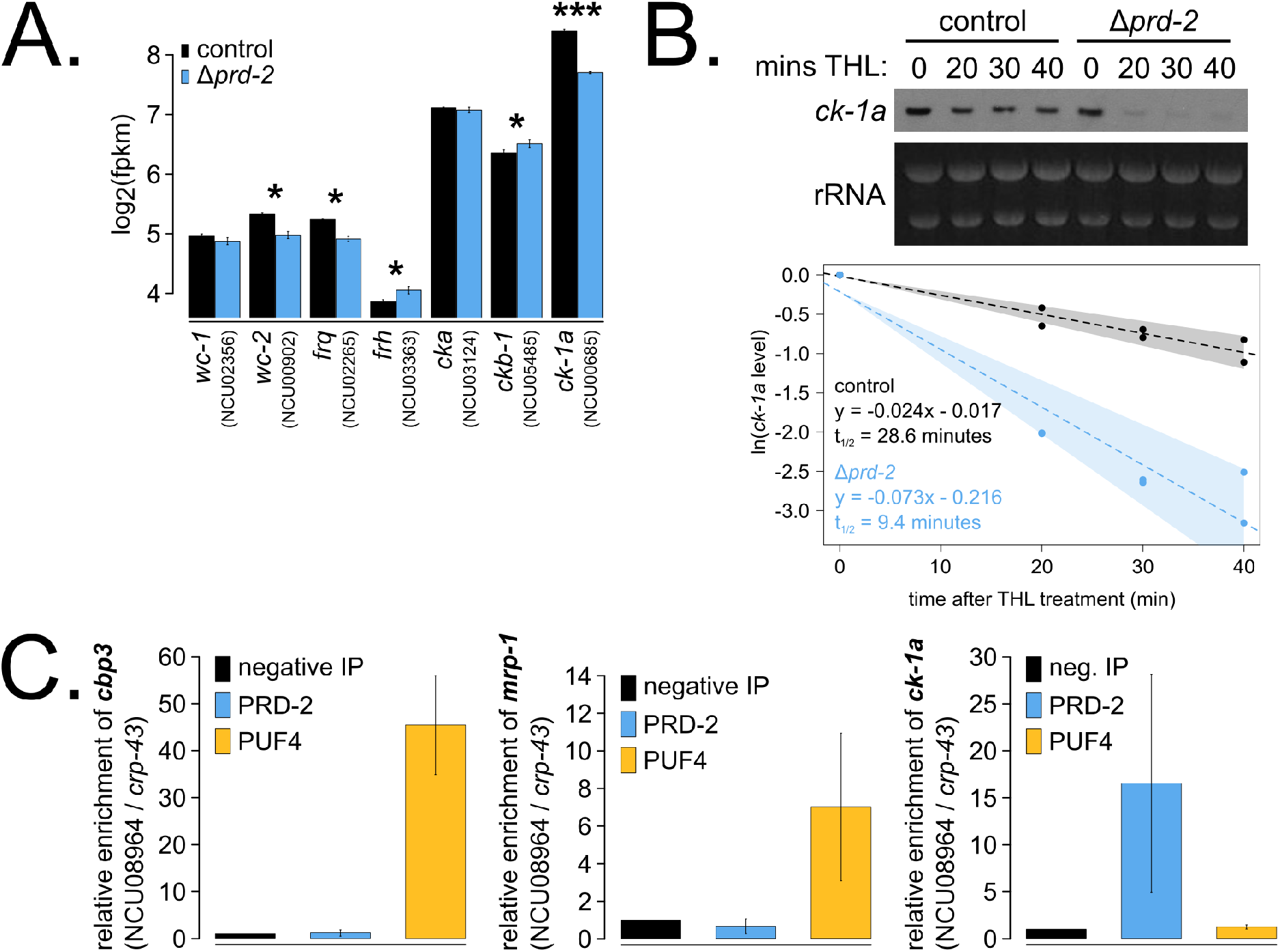
The core clock target of PRD-2 is the *casein kinase I* transcript. Control and *Δprd-2* cultures were grown in the light at 25°C in Bird medium for 48 hours prior to RNA isolation. Expression levels for core clock genes were measured by RNA-Sequencing (N = 3 biological replicates per strain), and log_2_-transformed FPKM values are shown. Asterisks indicate p < 0.05 (*) or p < 5×10^−5^ (***) by student’s t-test compared to control levels. The *ck-1a* transcript is > 1.5x less abundant in *Δprd-2* (**A**). *ck-1a* mRNA degradation kinetics were examined by Northern blot in a time course after treatment with thiolutin (THL) at approximately CT1 (N = 2 biological replicates). RNA levels were quantified using ImageJ, natural log transformed, fit with a linear model (glm in R, Gaussian family defaults), and half-life was calculated assuming first order decay kinetics (ln(2) / slope). Shaded areas around the linear fit represent 95% confidence intervals on the slope. The *ck-1a* transcript is 3x less stable in *Δprd-2* (**B**). The PUF4 (NCU16560) RNA-binding protein pulls down known target transcripts *cbp3* (NCU00057) and *mrp-1* (NCU07386) by RT-qPCR (N = 3 biological replicates). PRD-2 CLIP samples were processed in parallel with PUF4 positive controls, and PRD-2 binds the *ck-1a* transcript *in vivo* (**C**).

Hypothesizing that the clock-relevant target of PRD-2 could be CKI, we used two genetic approaches to manipulate CKI activity in an attempt to rescue the *Δprd-2* long period phenotype. First, we placed the *ck-1a* gene under the control of the quinic acid inducible promoter (Mehra et al., 2009) and crossed this construct into the *Δprd-2* background. We found that increasing expression of *ck-1a* using high levels (10^−1^ – 10^−2^ M) of QA partially rescued the *Δprd-2* long period phenotype (Figure 4A). We also noticed a synergistic poor growth defect in the double mutant at 10^−4^ M QA, consistent with low levels of *ck-1a* (an essential gene in *Neurospora:* (Gorl et al., 2001; He et al., 2006)). There are two explanations for the lack of full rescue to periods shorter than 25 hours in the *Pqa-2-ck-1a Δprd-2* double mutant: 1) even at saturating 10^−1^ M QA induction, the *qa-2* promoter may not reach endogenous levels of *ck-1a* achieved under its native promoter, and/or 2) because PRD-2 acts directly as an RNA-binding protein for CKI transcripts, simply increasing levels of *ck-1a* RNA cannot fully rescue PRD-2’s role in stabilizing or positioning CKI transcripts in the cytoplasm.

**Figure 4.**
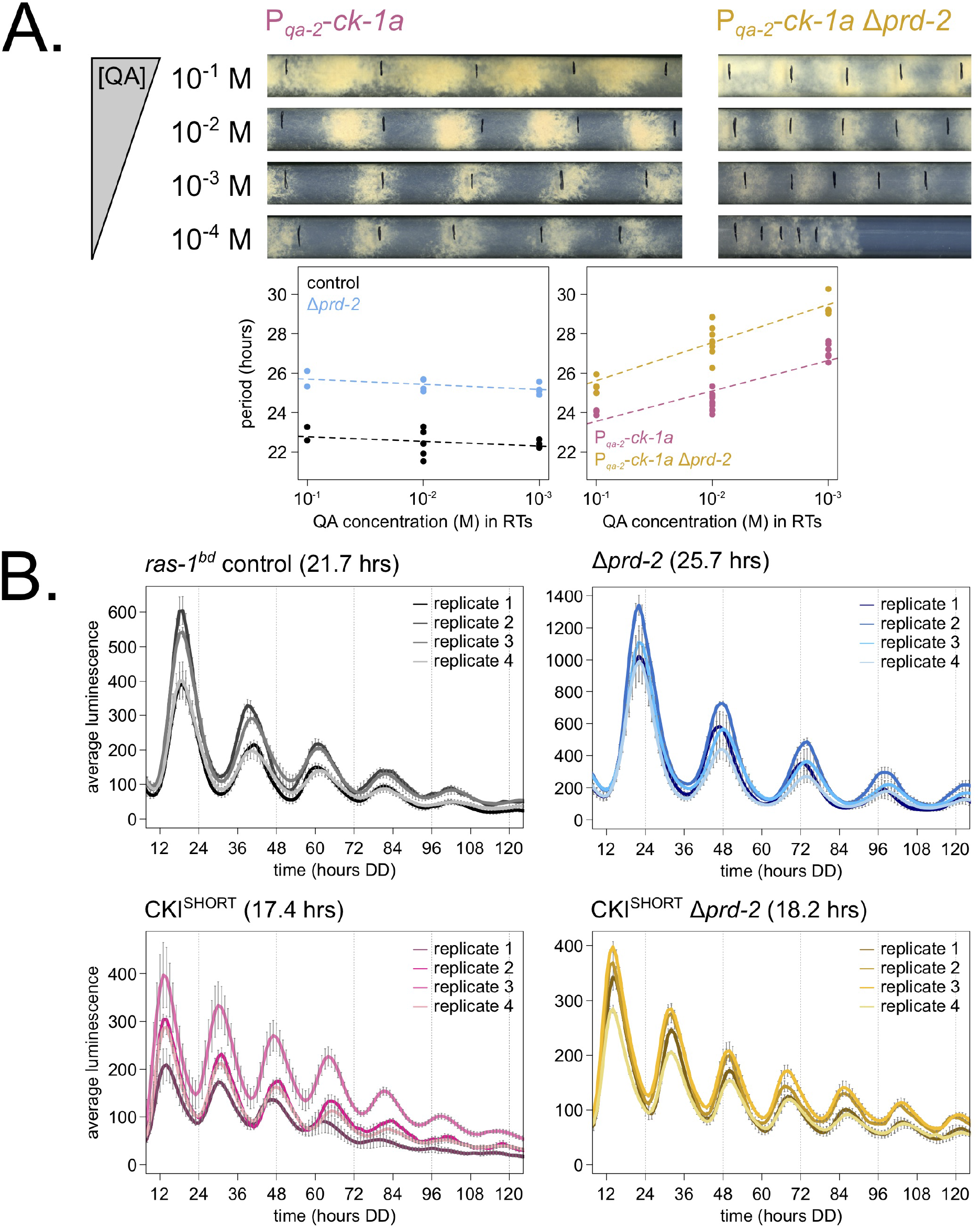
Genetically increasing CKI levels or activity rescues the *Δprd-2* long period phenotype. Representative race tubes from *ras-1^bd^ Pqa-2-ck-1a* single (pink) and *ras-1^bd^ Pqa-2-ck-1a Δprd-2* double (yellow) mutants are shown with growth using the indicated concentrations of quinic acid (QA) to drive expression of *ck-1a.* All results are shown in a scatterplot, where each dot represents one race tube’s free running period length. *ras-1^bd^* controls (black) had an average period of 22.5 ± 0.5 hours (N = 12), and period length was not significantly affected by QA concentration (ANOVA p = 0.297). *ras-1^bd^ Δprd-2* controls (blue) had an average period of 25.4 ± 0.4 hours (N = 10), and period length was not significantly affected by QA concentration (ANOVA p = 0.093). Period length of *ras-1^bd^ Pqa-2-ck-1a* single mutants (pink) was significantly altered across QA levels (ANOVA p = 3.6×10^−6^), and the average period at 10^−1^ M QA was 24.3 ± 0.5 hours (N = 4). Period length of *ras-1^bd^ Pqa-2-ck-1a Δprd-2* double mutants (yellow) was also significantly affected by QA levels (ANOVA p = 8.1×10^−8^), and the average period at 10^−1^ M QA was 25.4 ± 0.4 hours (N = 4). The double mutant period length was not genetically additive at high levels of QA induction (**A**). A hyperactive CKI allele was constructed by expressing the shortest isoform only (CKI^SHORT^). 96-well plate luciferase assays were used to measure the circadian period length. Traces represent the average of 3 technical replicates across 4 biological replicate experiments for: *ras-1^bd^* controls (gray, τ = 21.7 ± 0.3 hours), *ras-1^bd^ Δprd-2* (blue, τ = 25.7 ± 0.6 hours), *ras-1^bd^* CKI^SHORT^ (pink, τ = 17.4 ± 0.3 hours), and *ras-1^bd^* CKI^SHORT^ *Δprd-2* double mutants (yellow, τ = 18.2 ± 0.3). CKI^SHORT^ is completely epistatic to *Δprd-2* in double mutants (**B**).

Next, we turned to a previously described fungal CKI constitutively active allele, CKI Q299^STOP^ (Querfurth et al., 2007), reasoning that we might be able to rescue low *ck-1a* levels in *Δprd-2* by genetically increasing CKI kinase activity. We replaced endogenous CKI with a CKI^SHORT^ allele, which expresses only the shortest *ck-1a* isoform (361 amino acids). CKI^SHORT^ lacks 23 amino acids in the C-terminal tail of the full length isoform that are normally subject to autophosphorylation leading to kinase inhibition. This CKI^SHORT^ allele also carries an in-frame C-terminal HA3 tag and selectable marker, which displace the endogenous 3’ UTR of *ck-1a.* The CKI^SHORT^ mutant has a short period phenotype (~17 hrs), presumably due to hyperactive kinase activity and rapid feedback loop closure (Liu et al., 2019). Significantly, the CKI^SHORT^ mutation is completely epistatic to *Δprd-2* (Figure 4B), indicating that CKI is the clock-relevant target of PRD-2.

### Nonsense Mediated Decay Impacts the Clock by Regulating CKI Levels

Nonsense Mediated Decay (NMD) in *Neurospora crassa* is triggered by two different types of RNA structures. Open reading frames in 5’ UTRs that produce short peptides (5’ uORFs) can trigger NMD in a mechanism that does not require the Exon Junction Complex (Zhang and Sachs, 2015). The *frq* transcript has 6 such uORFs (Colot et al., 2005; Diernfellner et al., 2005) and could be a bona fide NMD target because its splicing is disrupted in the absence of NMD (Wu et al., 2017). In addition, transcripts with long 3’ UTRs, with intron(s) near a STOP codon, and/or with intron(s) in the 3’ UTR, can also be degraded by NMD after recruitment of the UPF1/2/3 complex by the Exon Junction Complex in a pioneering round of translation (Zhang and Sachs, 2015).

Since the observation by Compton (Compton, 2003) that the short period mutant *period-* 6 identified the UPF1 core subunit of the NMD pathway, the clock-relevant target(s) of NMD has been an object of conjecture and active research. Because loss of NMD reduces the amount of the transcript encoding the short-FRQ protein isoform (Wu et al., 2017), and strains making only short-FRQ have slightly lengthened periods (Liu et al., 1997), Wu et al. (2017) recently speculated that the short period of the *period-6^UPF1^* mutant might be explained by effects of NMD on FRQ. However, strains expressing only long-FRQ display an essentially wild-type period length (Colot et al., 2005; Liu et al., 1997), not a short period phenotype like *prd-6^UPF1^;* this finding is not consistent with FRQ being the only or even principal clock-relevant target of NMD, leaving unresolved the role of NMD in the clock.

To tackle this puzzle, we returned to classical genetic epistasis experiments and confirmed the observation that *prd-6* is completely epistatic to *prd-2^INV^* (Morgan and Feldman, 2001), going on to show that in fact each of the individual NMD subunit knockouts, *△upf2* and *△upf3* as well as *Δprd-6^upf1^*, is epistatic to the *Δprd-2* long period phenotype (Figure 5A). Previous work had profiled the transcriptome of *Δprd-6* compared to a control (Wu et al., 2017); we re-processed this RNA-Seq data and found, exactly as in *Δprd-2*, that *ck-1a* was the most affected core clock gene in *Δprd-6* (Figure 5B). The *ck-1a* transcript has an intron located 70 nt away from its longest isoform’s STOP codon, and its 3’ UTR is, remarkably, among the 100 longest annotated UTRs in the entire *Neurospora* transcriptome (Figure 5C). Thus, *ck-1a* is a strong candidate for regulation by NMD.

**Figure 5.**
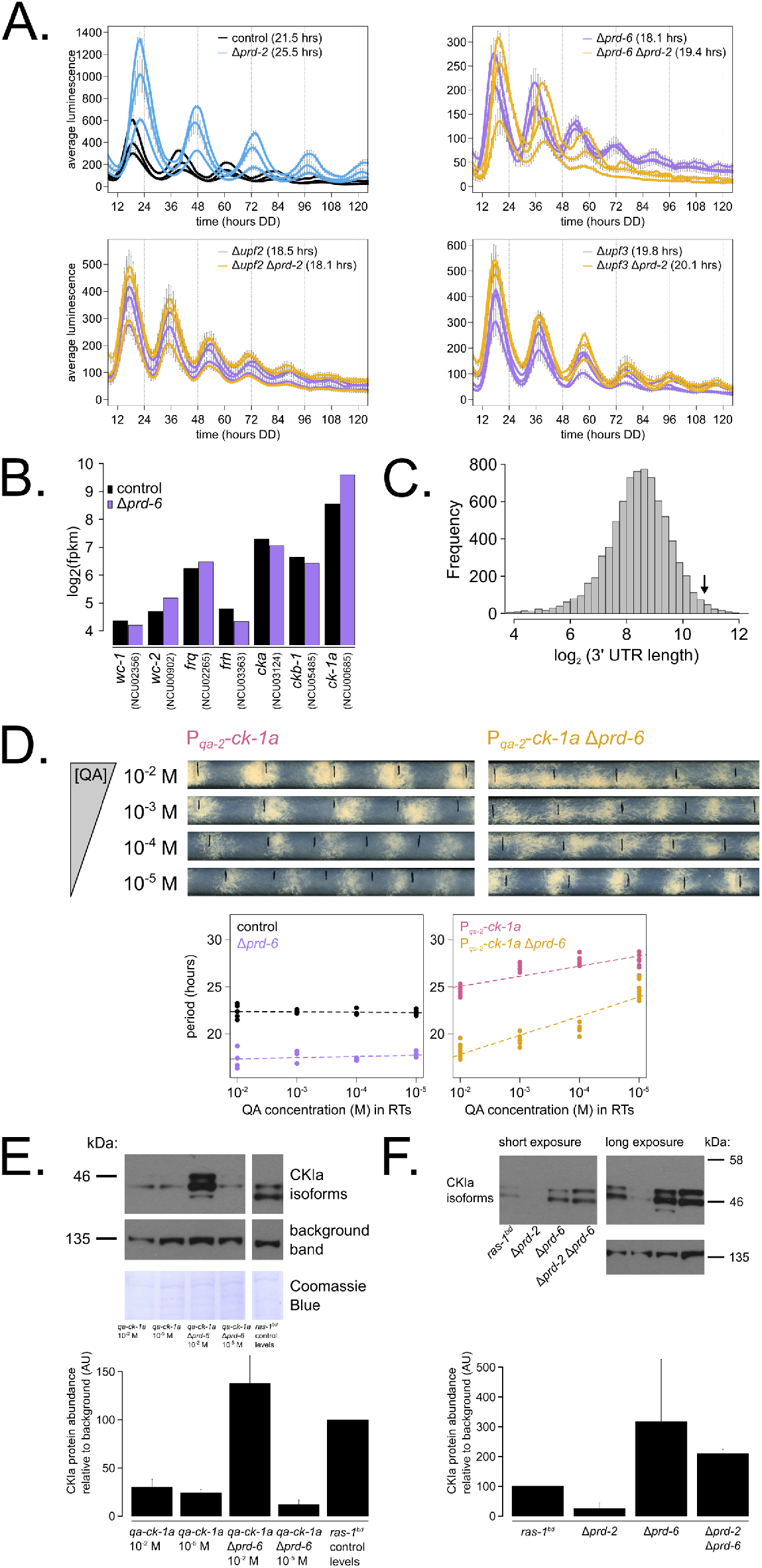
Nonsense Mediated Decay negatively regulates CKI levels via PRD-6, establishing a basis for the *prd-6 prd-2* genetic epistasis on circadian period length. 96-well plate luciferase assays were used to measure the circadian period length in triplicate wells per three biological replicate experiments for: *ras-1^bd^* controls (black, τ = 21.5 ± 0.3 hours), *ras-1^bd^ Δprd-2* (blue, τ = 25.5 ± 0.4 hours); *ras-1^bd^ Δprd-6* (purple, τ = 18.1 ± 0.2 hours), *ras-1^bd^ Δprd-6 Δprd-2* double mutants (yellow, τ = 19.4 ± 0.7 hours); *ras-1^bd^ Δupf2* (purple, τ = 18.5 ± 0.5 hours), *ras-1^bd^ Δupf2 Δprd-2* double mutants (yellow, τ = 18.1 ± 0.3 hours); *ras-1^bd^ Δupf3* (purple, τ = 19.8 ± 0.3 hours), *ras-1^bd^ Δupf3 Δprd-2* double mutants (yellow, τ = 20.1 ± 0.2 hours). Each individual NMD subunit knockout is epistatic to the *Δprd-2* long period phenotype (**A**). Raw RNA-Seq data from a previous study (Wu et al., 2017) were analyzed using the same pipeline as data from Figure 3A (see Materials & Methods). Control and *Δprd-6* gene expression levels (log_2_-transformed) are shown for core clock genes. The *ck-1a* transcript is > 2x more abundant in *Δprd-6* (**B**). 3’ UTR lengths from 7,793 genes were mined from the *N. crassa* OR74A genome annotation (FungiDB version 45, accessed on 10/25/2019), and plotted as a histogram. The arrow marks the 3’ UTR of *ck-1a*, which is 1,739 bp and within the top 100 longest annotated UTRs in the entire genome (**C**). Representative race tubes from *ras-1^bd^ Pqa-2-ck-1a* single (pink) and *ras-1^bd^ Pqa-2-ck-1a Δprd-6* double (yellow) mutants are shown at the indicated concentrations of quinic acid to drive expression of *ck-1a.* All results are shown in a scatterplot, where each dot represents one race tube’s free running period length. *ras-1^bd^* controls (black) had an average period of 22.4 ± 0.4 hours (N = 20), and period length was not significantly affected by QA concentration (ANOVA p = 0.605). *ras-1^bd^ Δprd-6* controls (purple) had an average period of 17.5 ± 0.6 hours (N = 16), and period length was not significantly affected by QA concentration (ANOVA p = 0.362). Period length of *ras-1^bd^ Pqa-2-ck-1a* single mutants (pink) was significantly altered across QA levels (ANOVA p = 2.9×10^−8^), and the average period at 10^−5^ M QA was 27.6 ± 0.8 hours (N = 8). Period length of *ras-1^bd^ Pqa-2-ck-1a Δprd-6* double mutants (yellow) was also significantly affected by QA levels (ANOVA p = 9.4×10^−12^), and the average period at 10^−5^ M QA was 24.7 ± 0.9 hours (N = 8). Thus, the double mutant period length was not genetically additive at low levels of QA induction, and the short period phenotype of *Δprd-6* is rescued (**D**). CKI protein levels were measured from the indicated genotypes grown in 0.1% glucose LCM medium with QA supplemented at the indicated concentrations for 48 hours in constant light. A representative immunoblot of 3 biological replicates is shown, and replicates are quantified in the bar graph relative to *ras-1^bd^* control CKI levels from a 2% glucose LCM culture (**E**). CKI protein levels were measured from the indicated genotypes grown in 2% glucose LCM medium for 48 hours in constant light. A representative immunoblot of 3 biological replicates is shown, and replicates are quantified in the bar graph relative to *ras-1^bd^* control CKI levels (**F**). CKI protein levels are increased in *Δprd-6*, decreased in the *Δprd-2* mutant, and *Δprd-6* is epistatic to *Δprd-2* with respect to CKI levels and circadian period length.

We hypothesized that CKI is overexpressed in the absence of NMD (Figure 5B), leading to faster feedback loop closure and a short circadian period. To genetically control *ck-1a* levels, we crossed the regulatable *Pqa-2-ck-1a* allele into the *Δprd-6* background and confirmed our hypothesis by finding that at low levels of inducer (10^−5^ M QA), decreased levels of *ck-1a* transcript revert the short period length of *Δprd-6* to control period lengths (Figure 5D). Further, protein levels of CKI in the *Δprd-6* background are reduced to control levels at 10^−5^ M QA (Figure 5E), which explains the period rescue phenotype. CKI protein is 2-3x more abundant in *Δprd-6* and in *Δprd-2 Δprd-6* (Figure 5F), matching its overexpression in the *Δprd-6* transcriptome (Figure 5B). CKI protein is 3x reduced in *Δprd-2* (Figure 5F), also correlating with its reduced mRNA expression and stability (Figure 3). We conclude that CKI is also the clock-relevant target of PRD-6^UPF1^, placing NMD, PRD-2, and CKI in the same genetic epistasis pathway.

## Discussion

By uncovering the identity and mode of action of PRD-2 and exploring the mechanism of two classical *period* mutants, *prd-2* and *prd-6*, we found a common basis in regulation of CKI levels, which are under tight control in the *Neurospora* clock (Figure 6). That the mechanistic basis of action of two independently derived non-targeted clock mutants centers on regulation of the activity of a single enzyme, CKI, via two distinct mechanisms is noteworthy. Period-2 encodes an RNA-binding protein (Figures 1-2) that stabilizes the CKI transcript (Figure 3B). We demonstrate that CKI is the most important core clock target of PRD-2 by rescuing its long period mutant phenotype with a hyperactive CKI allele (Figure 4B). The predominantly cytoplasmic localization of PRD-2 (Figure 2D) is consistent with its action in protecting *ck-1a* transcripts from NMD and rounds out the model. PTBP1, an RNA-binding protein, protects its target transcripts from NMD-mediated degradation by binding in the 3’ UTR and blocking NMD recruitment in mouse (Ge et al., 2016), and future work will determine if PRD-2 functions similarly to PTBP1.

**Figure 6.**
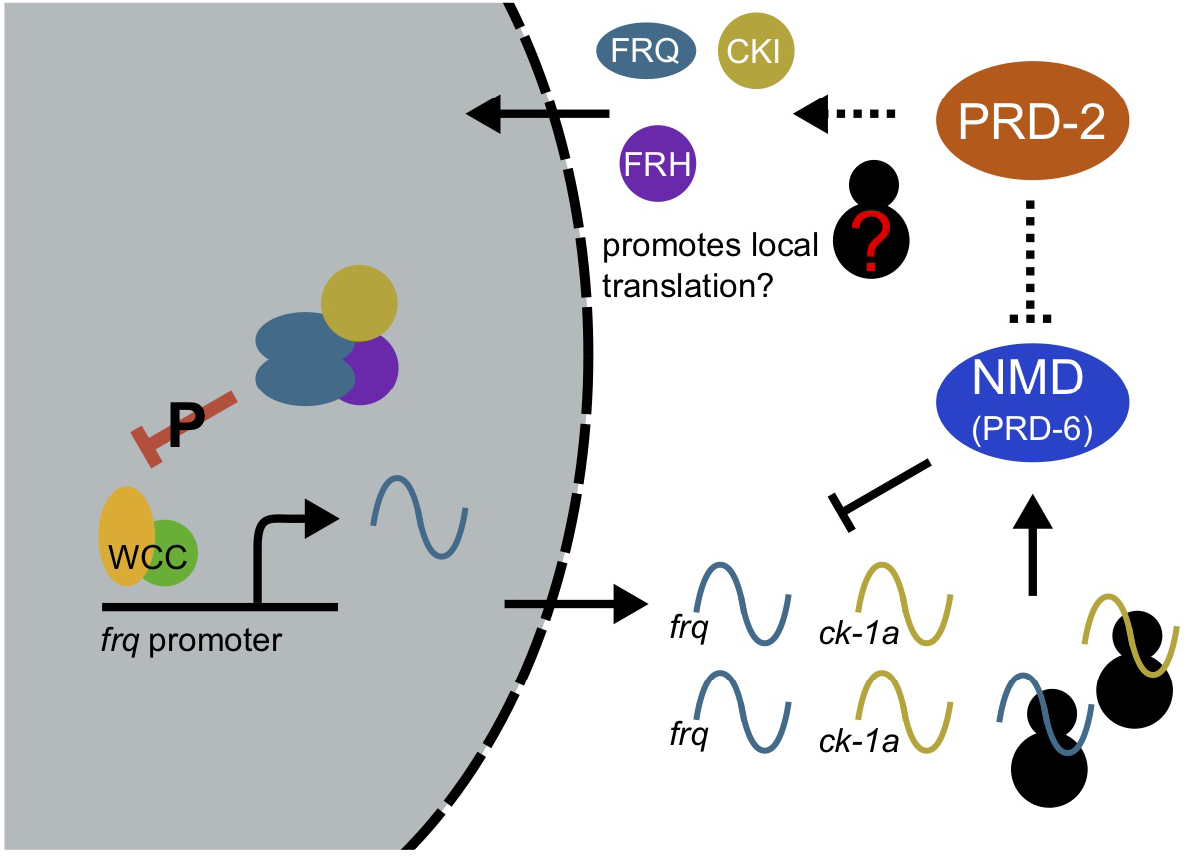
Counter balancing regulation of CKI provides a unifying genetic model for the action of PRD-2 and PRD-6^UPF-1^ in the circadian oscillator. The NMD complex (PRD-6^UPF1^, UPF2, and UPF3) targets the *frq* and *ck-1a* transcripts for degradation after the first round of translation (upstream uORFs in *frq;* long 3’ UTR in *ck-1a).* PRD-2 binds to and stabilizes *ck-1a* transcripts (dashed lines), which could also promote local translation and complex formation for the negative arm of the clock. In the absence of PRD-2, the long period phenotype is due to low CKI levels, and in the absence of NMD, the short period phenotype is due to high CKI levels.

PRD-6 and the NMD machinery target *ck-1a* mRNA for degradation to regulate its expression levels, presumably mediated by the long 3’ UTR of *ck-1a* transcripts in *Neurospora* (Figure 5). NMD components are not rhythmic in abundance in the fungal clock (Hurley et al., 2014, 2018). These data, taken together with the constitutive expression of the CKI mRNA and protein (Baker et al., 2009; Gorl et al., 2001; Hurley et al., 2014, 2018), lead us to predict that NMD regulation of CKI occurs throughout the circadian cycle. To our knowledge the discovery of NMD regulation of CKI represents a wholly novel and potentially important mode of regulation for this pivotal kinase. Future work will investigate whether insect DBT and/or mammalian CKIδ/ε (CSNK1D, CSNK1E) are also targets of NMD. Long UTR length appears to be conserved across CKI orthologs (Supplementary Figure 6). One previous study in *Drosophila* reported a circadian period defect in a tissue-specific NMD knockdown (Ri et al., 2019), but the behavioral rhythm was lengthened in UPF1-depleted insects unlike the short period defect observed in *Neurospora.* In mouse, both CKIε and CLOCK display altered splicing patterns in the absence of UPF2 (Weischenfeldt et al., 2012). Most core clock proteins have at least one uORF in mammals (Millius and Ueda, 2017), altogether raising the possibility that multiple core clock genes are regulated by NMD. The importance of NMD has already been recognized and investigated in the plant clock, where alternative splicing leads to NMD turnover for 4 core clock and accessory mRNAs: GRP7, GRP8, TOC1, and ELF3 (reviewed in: (Mateos et al., 2018)).

CKI abundance and alternative isoforms strongly affect circadian period length. Low levels of CKI driven from an inducible promoter lead to long periods approaching 30 hours (Mehra et al., 2009) (Figure 4A). In the mammalian clock, decreased CKI expression also significantly lengthens period (Isojima et al., 2009; Lee et al., 2009; Tsuchiya et al., 2016). CKI is rendered hyperactive by removing its conserved C-terminal domain, a domain normally subject to autophosphorylation leading to kinase inhibition (Guo et al., 2019; Querfurth et al., 2007). We generated a CKI mutant expressing only this shortest CKI isoform, finding a 17.5 hour short period phenotype in the absence of C-terminal autophosphorylation (Figure 4B). Based on prior work, increased CKI activity and/or abundance would be expected to increase FRQ-CKI affinity and lead to faster feedback loop closure (Liu et al., 2019), consistent with the short period phenotype. Curiously, this CKI short isoform is expressed at levels similar to the full length isoform in *Neurospora* (as well as a third short isoform derived from an alternative splice acceptor event) (Figure 5F), and all isoforms interact with FRQ by immunoprecipitation (Querfurth et al., 2007). Why do natural isoforms arise without the auto-inhibitory C-terminus in *Neurospora*, and are these regulatory events required to keep the clock on time? Mammalian alternative isoforms *CKIδ1 and CKIδ2* have different substrate preferences *in vitro*, which leads to differential phosphorylation of PER2 whereby CKIδ2 phosphorylation significantly stabilizes PER2 (Fustin et al., 2018). Adding further complexity, *CKIδ1 and CKIδ2* isoform expression patterns appear to be tissue specific and are regulated by m6A RNA modification. Regulation of CKI levels and isoform expression is an important direction for future work in the circadian clock.

Casein Kinase I has a diverse array of functions in eukaryotes and is critically important in human health (reviewed in: (Cheong and Virshup, 2011; Vielhaber and Virshup, 2001)). CKI overexpression is pathogenic in Alzheimer’s Disease in addition to its role in circadian period regulation (Sundaram et al., 2019). Future work on regulation of CKI levels and isoform expression will shed light on CKI regulation in the clock, in development, and in disease.

## Materials and Methods

### *Neurospora* strains and growth conditions

The *ras-1^bd^ prd-2^INV^* strains 613-102 (*mat* A) and 613-43 (*mat* a) were originally isolated in the Feldman laboratory (Lewis, 1995). Strains used in this study were derived from the wild-type background (FGSC2489 *mat* A), *ras-1^bd^* background (87-3 *mat* a or 328-4 *mat* A), *Δmus-51* background (FGSC9718 *mat* a), or the Fungal Genetics Stock Center (FGSC) knockout collection as indicated (Supplementary Table 1). Strains were constructed by transformation or by sexual crosses using standard *Neurospora* methods (http://www.fgsc.net/Neurospora/NeurosporaProtocolGuide.htm). In the “c box-luc” core clock transcriptional reporter used throughout, a codon-optimized firefly luciferase gene is driven by the clock box in the *frequency* promoter (Gooch et al., 2008; Hurley et al., 2014; Larrondo et al., 2015). The clock reporter construct was targeted to the *csr-1* locus and selected on resistance to 5 μg/ml cyclosporine A (Sigma # 30024) (Bardiya and Shiu, 2007).

Standard race tube (RT) medium was used for all race tubes (1X Vogel’s Salts, 0.1% glucose, 0.17% arginine, 1.5% agar, and 50 ng/ml biotin). Where indicated, D-Quinic Acid (Sigma # 138622) was added from a fresh 1 M stock solution (pH 5.8). Standard 96-well plate medium was used for all camera runs (1X Vogel’s Salts, 0.03% glucose, 0.05% arginine, 1.5% agar, 50 ng/ml biotin, and 25 μM luciferin from GoldBio # 115144-35-9). Liquid cultures were started from fungal plugs as described (Chen et al., 2009; Nakashima, 1981) or from a conidial suspension at 1×10^5^ conidia/ml. Liquid cultures were grown in 2% glucose Liquid Culture Medium (LCM) or in 1.8% glucose Bird Medium (Metzenberg, 2004) as indicated. QA induction experiments in liquid culture were performed in 0.1% glucose LCM medium with QA supplemented. All experiments were conducted at 25°C in constant light unless otherwise indicated.

Strains were genotyped by screening for growth on selection medium (5 μg/ml cyclosporine A, 400 μg/ml Ignite, and/or 200-300 μg/ml Hygromycin). PCR genotyping was performed on gDNA extracts from conidia incubated with Allele-In-One Mouse Tail Direct Lysis Buffer (Allele Biotechnology # ABP-PP-MT01500) according to the manufacturer’s instructions. GreenTaq PCR Master Mix (ThermoFisher # K1082) was used for genotyping. Relevant genotyping primers for key strains are:

*ras-1^bd^* (mutant): 5’ TGCGCGAGCAGTACATGCGAAT and 5’

CCTGATTTCGCGGACGAGATCGTA 3’

*ras-1^WT^* (NCU08823): 5’ GCGCGAGCAGTACATGCGGAC 3’ and 5’

CCTGATTTCGCGGACGAGATCGTA 3’

*prd-2^WT^* (NCU01019): 5’ CACTTCCAGTTATCTCGTCAC 3’ and 5’

CACAACCTTGTTAGGCATCG 3’

Δ*prd-2*::bar^R^ (KO mutant): 5’ CACTTCCAGTTATCTCGTCAC 3’ and 5’

GTGCTTGTCTCGATGTAGTG 3’

*prd-2^INV^* (left breakpoint): 5’ AGCGAGCTGATATGCCTTGT 3’ and 5’

CGACTTCCACCACTTCCAGT 3’

*prd-2^INV^* (right breakpoint): 5’ TGTTTGTCCGGTGAAGATCA 3’ and 5’

GTCGTGGAATGGGAAGACAT 3’

Δ*prd-6*::hyg^R^ (FGSC KO mutant): 5’ CTGCAACCTCGGCCTCCT 3’ and 5’ CAGGCTCTCGATGAGCTGATG 3’

bar^R^::P*qa-2*-*ck-1a* (QA inducible CKI): 5’ GTGCTTGTCTCGATGTAGTG 3’ and 5’ GATGTCGCGGTGGATGAACG 3’

### RNA stability assays

Control and *Δprd-2* liquid cultures grown in 1.8% glucose Bird medium were age-matched and circadian time (CT) matched to ensure that RNA stability was examined at the same phase of the clock. Control cultures were shifted to constant dark for 12 hours, and *Δprd-2* cultures were shifted to dark for 14 hours (~CT 1 for 22.5-hour wild-type period and for 26-hour *Δprd-2* period; 46 hours total growth). Thiolutin (THL; Cayman Chemical # 11350) was then added to a final concentration of 12 μg/ml to inhibit new RNA synthesis. Samples were collected every 10 minutes after THL treatment by vacuum filtration and flash frozen in liquid nitrogen. THL has multiple off-target effects in addition to inhibiting transcription (Lauinger et al., 2017). For this reason, *frq* mRNA degradation kinetics were also examined with an alternative protocol. Light-grown, age-matched liquid Bird cultures of wild-type and *Δprd-2* were shifted into the dark and sampled every 10 minutes to measure *frq* turnover; transcription of *frq* ceases immediately on transfer to darkness (Heintzen et al., 2001; Tan et al., 2004). All tissue manipulation in the dark was performed under dim red lights, which do not reset the *Neurospora* clock (Chen et al., 2009).

### RNA isolation and detection

Frozen *Neurospora* tissue was ground in liquid nitrogen with a mortar and pestle. Total RNA was extracted with TRIzol (Invitrogen # 15596026) and processed as described (Chen et al., 2009). RNA samples were prepared for RT-qPCR, Northern Blotting, RNA-Sequencing, or stored at −80°C.

For RT-qPCR, cDNA was synthesized using the SuperScript III First-Strand synthesis kit (Invitrogen # 18080-051). RT-qPCR was performed using SYBR green master mix (Qiagen # 204054) and a StepOne Plus Real-Time PCR System (Applied Biosystems). Ct values were determined using StepOne software (Life Technologies) and normalized to the *actin* gene (ΔC_t_). The ΔΔC_t_ method was used to determine mRNA levels relative to a reference time point. Relevant RT-qPCR primer sequences are: *prd-2* (NCU01019): 5’ GGGCAACGACGTCAAACTAT 3’ and 5’ TGCGTGTACATCACTCTGGA 3’. *actin* (NCU04173): 5’ GGCCGTGATCTTACCGACTA 3’ and 5’ TCTCCTTGATGTCACGAACG 3’.

Northern probes were first synthesized using the PCR DIG Probe Synthesis Kit (Roche # 11 636 090 910). The 512 bp *frq* probe was amplified from wild-type *Neurospora* genomic DNA with primers: 5’ CTCTGCCTCCTCGCAGTCA 3’ and 5’ CGAGGATGAGACGTCCTCCATCGAAC 3’. The 518 bp *ck-1a* probe was amplified with primers: 5’ CCATGCCAAGTCGTTCATCC 3’ and 5’ CGGTCCAGTCAAAGACGTAGTC 3’. Total RNA samples were prepared according to the NorthernMax™-Gly Kit instructions (Invitrogen # AM1946). Equal amounts of total RNA (5 – 10 μg) were loaded per lane of a 0.8 – 1% w/v agarose gel. rRNA bands were visualized prior to transfer to validate RNA integrity. Transfer was completed as described in the NorthernMax™-Gly instructions onto a nucleic acid Amersham Hybond-N+ membrane (GE # RPN303B). Transferred RNA was crosslinked to the membrane using a Stratalinker UV Crosslinker. The membrane was blocked and then incubated overnight at 42°C in hybridization buffer plus the corresponding DIG probe. After washing with NorthernMax™-Gly Kit reagents, subsequent washes were performed using the DIG Wash and Block Buffer Set (Roche # 11 585 762 001). Anti-Digoxigenin-AP Fab fragments were used at 1:10,000. Chemiluminescent detection of anti-DIG was performed using CDP-Star reagents from the DIG Northern Starter Kit (Roche # 12 039 672 910). Densitometry was performed in ImageJ.

Total RNA was submitted to Novogene for stranded polyA+ library preparation and sequencing. 150 bp paired-end (PE) read libraries were prepared, multiplexed, and sequenced in accordance with standard Illumina HiSeq protocols. 24.8 ± 1.7 million reads were obtained for each sample. Raw FASTQ files were aligned to the *Neurospora crassa* OR74A NC12 genome (accessed September 28, 2017 via the Broad Institute: ftp://ftp.broadinstitute.org/pub/annotation/fungi/neurospora_crassa/assembly/) using STAR (Dobin et al., 2013). On average, 97.6 ± 0.3% of the reads mapped uniquely to the NC12 genome. Aligned reads were assembled into transcripts, quantified, and normalized using Cufflinks2 (Trapnell et al., 2013). Triplicate control and *Δprd-2* samples were normalized together with CuffNorm, and the resulting FPKM output was used in the analyses presented. RNA-Sequencing data have been submitted to the NCBI Gene Expression Omnibus (GEO; https://www.ncbi.nlm.nih.gov/geo/) under accession number GSE155999.

### CLIP assay

CLIP was performed using PUF4 (NCU16560) as a positive control RNA-binding protein from (Wilinski et al., 2017), with modifications. *Neurospora* strains containing endogenous locus C-terminally VHF tagged PUF4, PRD-2, or untagged negative control were used (Supplementary Table 1). Liquid cultures were grown in 2% glucose LCM for 48 hours in constant light. Tissue was harvested by vacuum filtration and fixed by UV crosslinking for 7 minutes on each side of the fungal mat (Stratalinker UV Crosslinker 1800 with 254-nm wavelength bulbs). UV crosslinked tissue was frozen in liquid nitrogen and ground into a fine powder with a mortar and pestle. Total protein was extracted in buffer (25 mM Tris-HCl pH 7.4, 150 mM NaCl, 2 mM MgCl2, 0.5% NP-40, 1 mM DTT, 1x cOmplete protease inhibitor, 100 U / ml RNAse Out) and concentration determined by Bradford Assay. Approximately 10 mg of total protein was added to 30 μl anti-FLAG M2 magnetic beads (Sigma # M8823) prepared according to the manufacturer’s instructions. Beads and lysate were rotated for 4 hours at 4°C, followed by 4 washes in 750 μl extraction buffer. Bound RNA-binding proteins were eluted with 100 μl 0.1 M glycine-HCl pH 3.0 for 10 minutes. The supernatant was collected using a magnetic rack (NEB S1506S) and neutralized in 10 μl of 1 M Tris pH 8.0. The elution was incubated with 300 μl of TRIzol (Invitrogen # 15596026) for 10 minutes to extract RNA. Total RNA was isolated, DNAse treated, and concentrated using the Direct-zol RNA Microprep Kit (Zymo # R2062) following the manufacturer’s instructions.

Equal amounts of immunoprecipitated RNA (~ 50 ng) were converted into cDNA using the oligo(dT) method from the SuperScript IV First-Strand synthesis kit (Invitrogen # 1809-1050). RT-qPCR was performed using SYBR green master mix (Qiagen # 204054) and a StepOne Plus Real-Time PCR System (Applied Biosystems). Ct values were determined using StepOne software (Life Technologies) and normalized to the *crp-43* gene (ΔCt) instead of the *actin* (NCU04173) gene because *actin* is a putative PUF4 target by HITS-CLIP (Wilinski et al., 2017). The ΔΔC_t_ method was used to determine target mRNA enrichment relative to the negative IP sample. Relevant RT-qPCR primer sequences were designed to flank introns: *cbp3* (NCU00057; PUF4 target): 5’ CGAGAAATTCGGCCTTCTCCC 3’ and 5’ GCCTGGTGGAAGAAGTGGT 3’. *mrp-1* (NCU07386; PUF4 target): 5’ TAGTAGGCACCGACTTTGAGCA 3’ and 5’ CGGGGACAGGTGGTCGAA 3’. *ck-1a* (NCU00685; PRD-2 target): 5’ CGCAAACATGACTACCATG 3’ and 5’ CTCTCCAGCTTGATGGCA 3’. *crp-43* (NCU08964; normalization control): 5’ CTGTCCGTACTCGTGACTCC 3’ and 5’ ACCATCGATGAGGAGCTTGC 3’.

### Protein isolation and detection

Frozen *Neurospora* tissue was ground in liquid nitrogen with a mortar and pestle. Total protein was extracted in buffer (50 mM HEPES pH 7.4, 137 mM NaCl, 10% glycerol v/v, 0.4% NP-40 v/v, and cOmplete Protease Inhibitor Tablet according to instructions for Roche # 11 836 170 001) and processed as described (Garceau et al., 1997). Protein concentrations were determined by Bradford Assay (Bio-Rad # 500-0006). For Western blots, equal amounts of total protein (10 – 30 μg) were loaded per lane into 4-12% Bis-Tris Bolt gels (Invitrogen # NW04125BOX). Western transfer was performed using an Invitrogen iBlot system (# IB21001) and PVDF transfer stack (# IB401001). Primary antibodies used for Western blotting were anti-V5 (1:3000, ThermoFisher # R960-25), anti-Tubulin alpha (1:10,000, Fitzgerald # 10R-T130a), or anti-CK1a (1:1000, rabbit raised). The secondary antibodies, goat anti-mouse or goat anti-rabbit HRP, were used at 1:5000 (Bio-Rad # 170-6516, # 170-6515). SuperSignal West Pico PLUS Chemiluminescent Substrate (ThermoFisher # 34578) or Femto Maximum Sensitivity Substrate (Thermo # 34095) was used for detection. Immunoblot quantification and normalization were performed in ImageJ.

Nuclear and cytosolic fractions were prepared as previously described (Hong et al., 2008). Approximately 10 μg of total protein from each fraction were loaded for immunoblotting. Primary antibodies for fraction controls were histone H3A (Fitzgerald) and γ-tubulin (Abcam). HRP-conjugated secondary antibodies (Bio-Rad) were used with SuperSignal West Pico ECL (Thermo) for detection.

### Luciferase reporter detection and data analysis

96-well plates were inoculated with conidial suspensions from strains of interest and entrained in 12 hour light:dark cycles for 2 days in a Percival incubator at 25°C. Temperature inside the Percival incubator was monitored using a HOBO logger device (Onset # MX2202) during entrainment and free run. Plates were then transferred into constant darkness to initiate the circadian free run. Luminescence was recorded using a Pixis 1024B CCD camera (Princeton Instruments). Light signal was acquired for 10 – 15 minutes every hour using LightField software (Princeton Instruments, 64-bit version 6.10.1). The average intensity of each well was determined using a custom ImageJ Macro (Larrondo et al., 2015), and background correction was performed for each frame. Results from two different algorithms were averaged together to determine circadian period from background-corrected luminescence traces. The MESA algorithm was used as previously described (Kelliher et al., 2020). A second period measurement was obtained from an ordinary least squares autoregressive model to compute the spectral density (in R: `spec.ar(…, method = “¤ls”`)). Race tube period lengths were measured from scans using ChronOSX 2.1 software (Roenneberg and Taylor, 2000).

### Data visualization

All figures were plotted in R, output as scalable vector graphics, formatted using Inkscape, and archived in R markdown format. Data represent the mean of at least three biological replicates with standard deviation error bars, unless otherwise indicated.

## Supplementary Figures

**Supplementary Figure 1.**
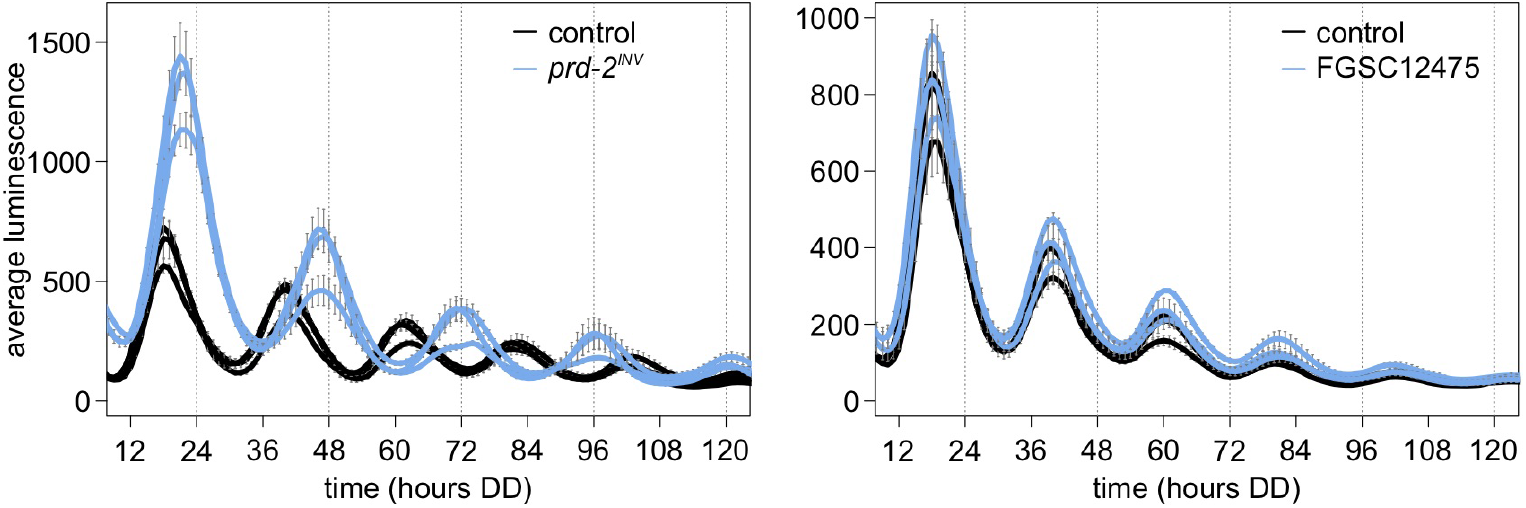
NCU03775 knockout has a normal circadian period and does not explain the *prd-2^INV^* phenotype. 96-well plate luciferase assays were used to measure the circadian period length. Traces represent the average of 3 technical replicates across 3 biological replicate experiments for: *ras-1^bd^* controls (black, τ = 21.9 ± 0.3 hours), *ras-1^bd^ prd-2^INV^* (blue, τ = 25.6 ± 0.4 hours), wild-type controls (black, τ = 21.7 ± 0.3 hours), and the knockout strain FGSC12475 (blue, τ = 21.7 ± 0.3 hours). ΔNCU03775 has a wild-type circadian period length.

**Supplementary Figure 2.**
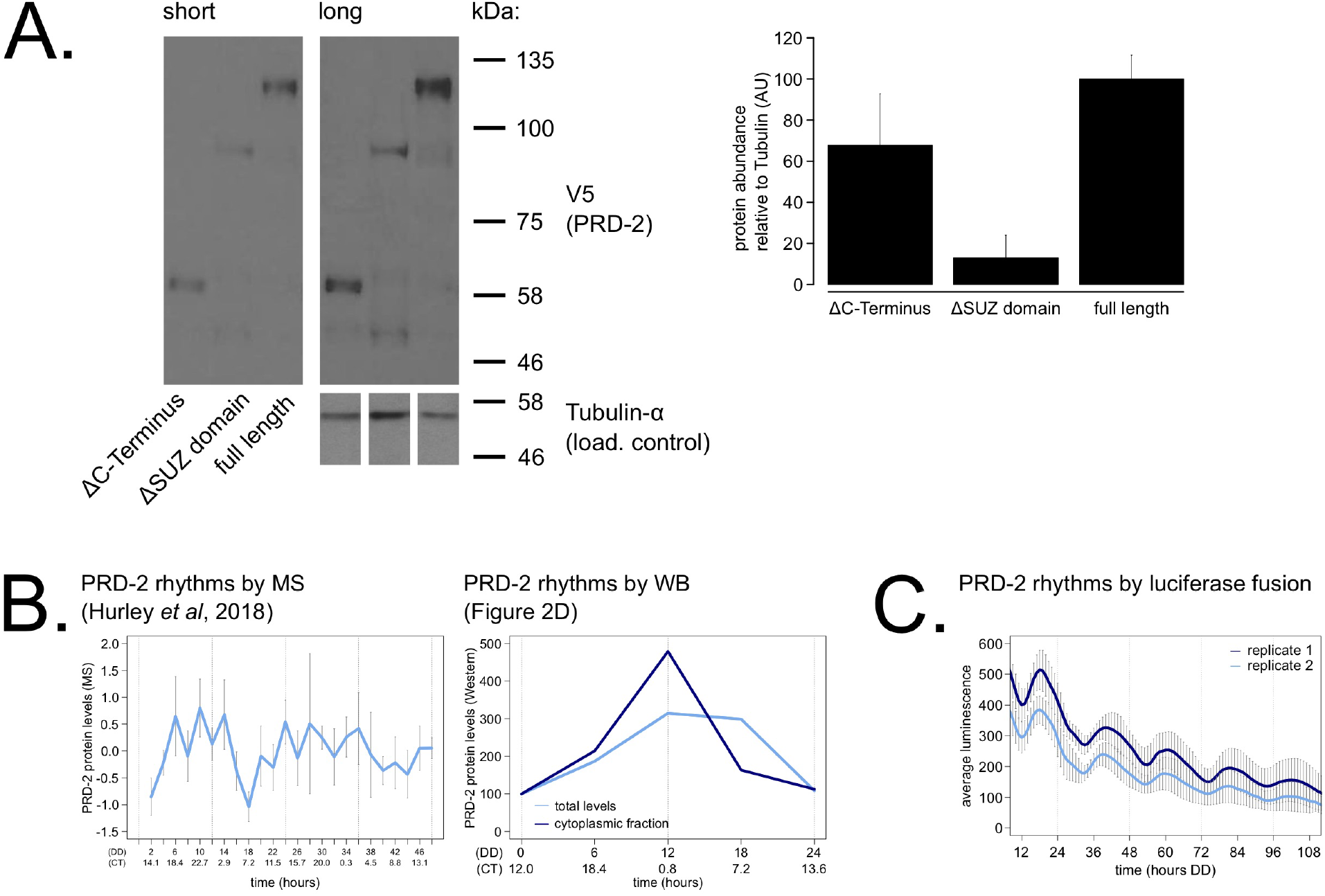
PRD-2 protein levels are slightly rhythmic and are detectable in protein domain deletion mutants. PRD-2 protein levels were measured from at least 3 biological replicates using strains where the endogenous NCU01019 locus was replaced with V5-tagged domain deletion constructs: *prd-2*ΔC-Terminus(Δ440-790), *prd-2* ΔSUZ(Δ345-431), and full length. Long and short exposures of a representative immunoblot are shown with quantification relative to tubulin loading controls (**A**). The C-Terminal deletion strain has ~68% PRD-2 levels compared to the full length control, and the SUZ deletion has ~13% levels. Both are above the low levels of *qa*-driven NCU01019 needed to induce the long period phenotype (Figure 1E). MS data from a previous study (Hurley et al., 2018) revealed low amplitude rhythms in PRD-2 abundance with a broad peak in the subjective circadian night and early morning (~ CT18 – 2). Circadian Time (CT) was calculated as described previously (Kelliher et al., 2020). PRD-2 abundance was also quantified from the localization time course (Figure 2D) relative to tubulin and relative to the first time point. Peak PRD-2 protein abundance was observed in the subjective morning from both MS and immunoblot data (**B**), corresponding with the rise in *frq* transcript levels (Aronson et al., 1994). To confirm rhythms in PRD-2 protein expression, the complete NCU01019 5’ UTR and coding sequence, plus 951 bp of upstream promoter sequence, was fused in-frame with codon-optimized luciferase (Gooch et al., 2008), not including its endogenous 3’ UTR sequence. This construct was transformed into the *prd-2*^WT^ background at the *csr-1* locus, and PRD-2 protein cycles in abundance (τ = 21.7 ± 0.8 hours). PRD-2 protein peaks during the circadian day (CT7.5 ± 1) by luciferase fusion (**C**), slightly delayed relative to its morning peak by Western and MS.

**Supplementary Figure 3.**
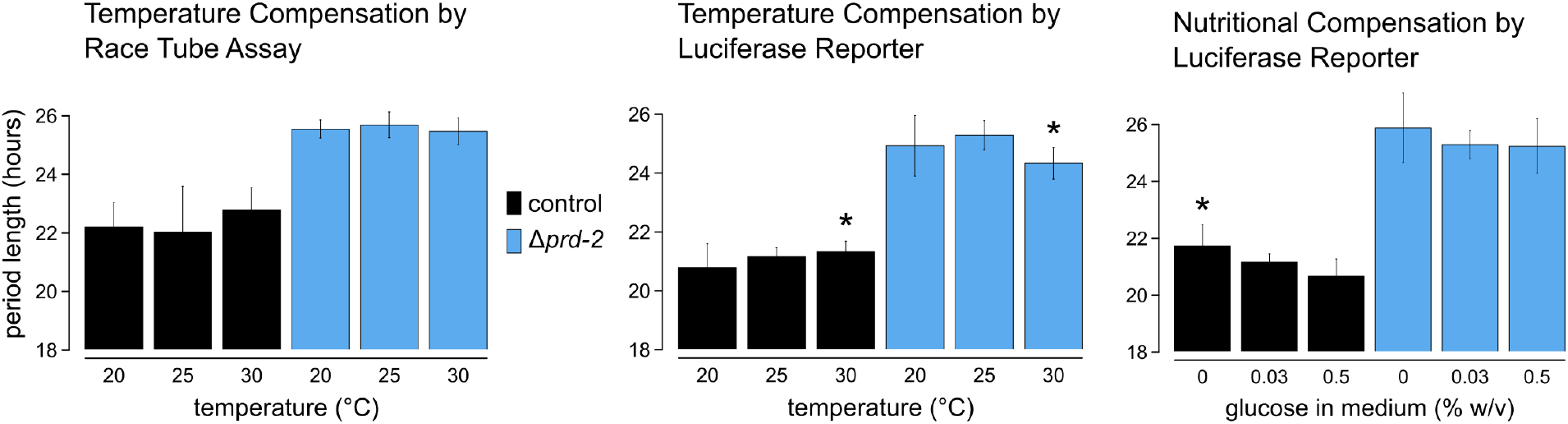
Temperature and nutritional compensation are normal in ΔNCU01019. Temperature and nutritional compensation were assessed in *ras-1^bd^* controls compared to *ras-1^bd^ Δprd-2.* Race tubes were incubated at 20°, 25°, or 30°C to determine free running period length. Temperature did not significantly affect period length for controls (ANOVA p = 0.598) or for the *Δprd-2* mutant (ANOVA p = 0.756) race tubes. 96-well plates were incubated at 20°, 25°, or 30°C to determine free running period length. Period was significantly different at 30°C for both genotypes (Asterisks (*): control 20°C vs 30°C, Tukey Test p = 0.023; *Δprd-2* 20°C vs 30°C, Tukey Test p = 0.037; *Δprd-2* 25°C vs 30°C, Tukey Test p = 0.0002). 96-well plates were run with 0%, 0.03%, or 0.5% glucose w/v to test nutritional compensation. Period length was significantly different at 0% glucose for controls only (Asterisk (*): control 0% vs 0.5%, Tukey Test p = 0.00005; control 0% vs 0.03%, Tukey Test p = 0.034; *Δprd-2* ANOVA p = 0.183).

**Supplementary Figure 4.**
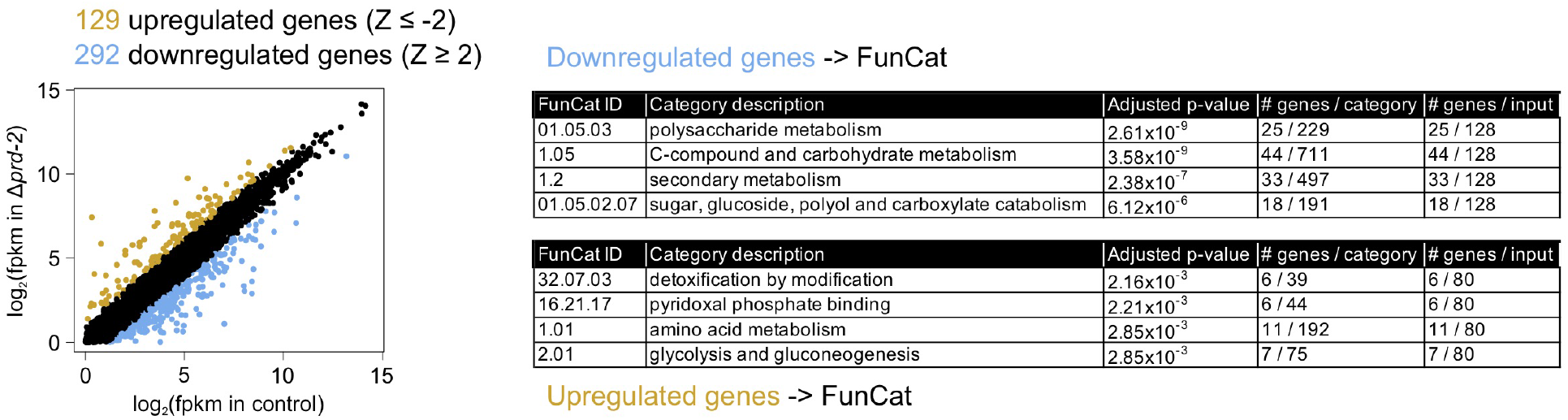
Hundreds of genes have altered expression levels in the *Δprd-2* mutant but a common pathway or sequence motif was not detected. RNA-Seq data were first filtered for low expression. 8,622 out of 9,730 annotated *N. crassa* genes were expressed in 4/6 samples (> 0 FPKM units in triplicate control and *Δprd-2).* FPKM units for 8,622 expressed genes were log_2_-transformed, averaged, subtracted from control, and Z-scores computed. 129 genes (gold) were upregulated in *Δprd-2* (Z-score < −2), and 292 genes (blue) were downregulated in *Δprd-2* (Z-score > 2). Hypothesizing that PRD-2 is an RNA-binding protein that stabilizes its target transcripts (Figure 3B), we searched for enriched sequence motifs in the untranslated regions of the 292 downregulated genes using Weeder2 (212 annotated 5’ UTRs and 226 annotated 3’ UTRs searched). Zero motifs scored better than 1.5 from Weeder2 output compared to background *Neurospora* nucleotide frequencies (data not shown). Up- and down-regulated gene categories were then run through FunCat to determine functionally enriched categories of genes in the putative PRD-2 regulon. 128 of the 292 downregulated genes were input to FunCat, and the top scoring functional categories indicated that carbohydrate and secondary metabolism were decreased in *Δprd-2.* 80 out of the 128 upregulated genes were also input to FunCat, and other metabolism categories were identified, which could indicate altered central carbon metabolism in the *Δprd-2* mutant, correlating with its slow growth phenotype.

**Supplementary Figure 5.**
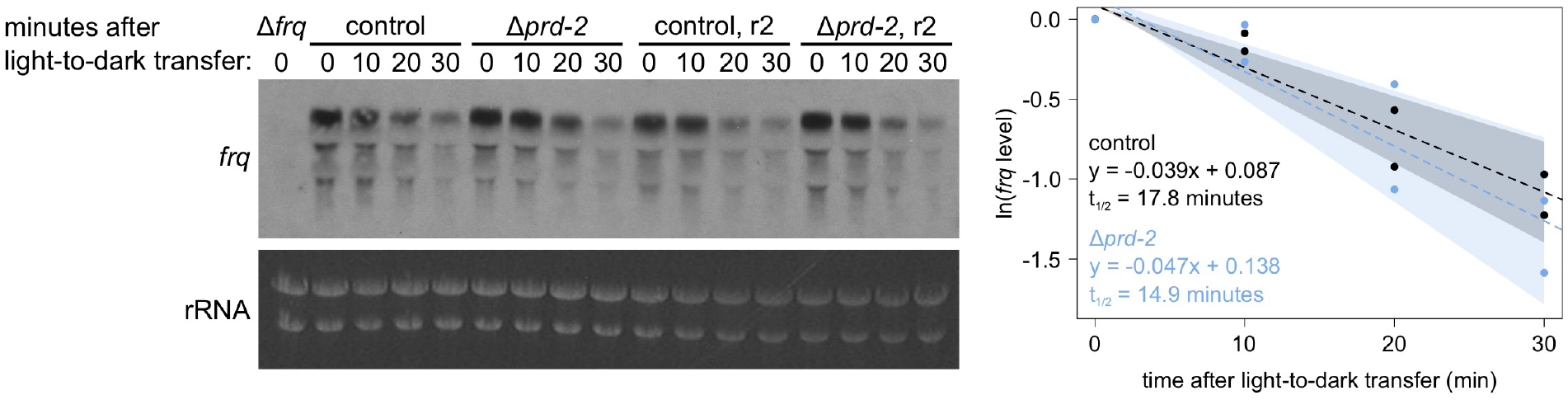
Loss of *prd-2* has little effect on stability of the *frq* transcript. *frq* mRNA degradation kinetics were examined by Northern blot in a time course after light-to-dark transfer (N = 2 biological replicates). RNA levels were quantified using ImageJ, natural log transformed, fit with a linear model (`glm` in R, Gaussian family defaults), and half-life was calculated assuming first order decay kinetics (ln(2) / slope). Shaded areas around the linear fit represent 95% confidence intervals on the slope. The *frq* half-life is approximately 3 minutes shorter in *Δprd-2* but is not statistically different from the control.

**Supplementary Figure 6.**
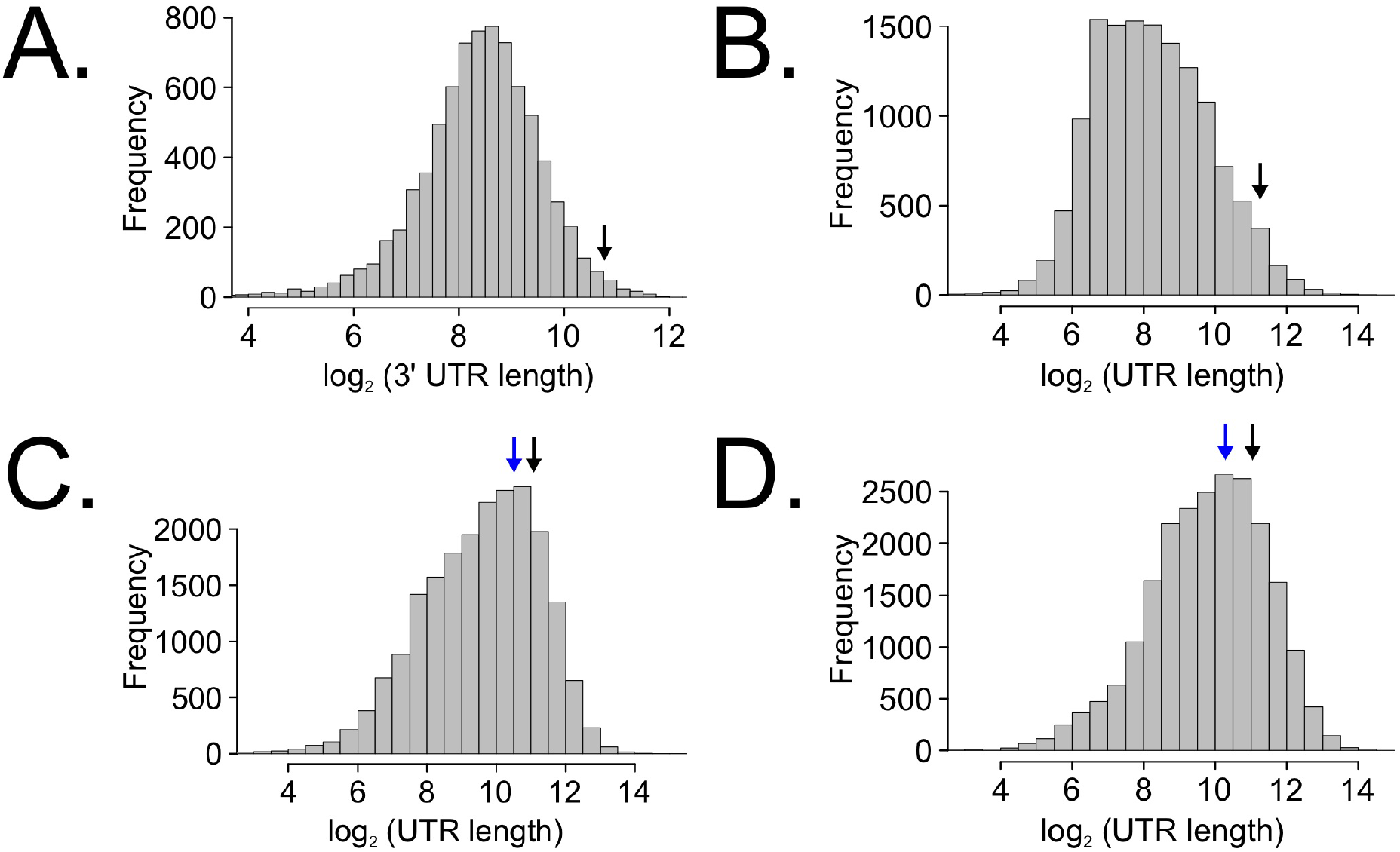
Long untranslated regions (UTRs) are characteristic of *casein kinase I* gene orthologs across species. *Neurospora* 3’ UTR lengths were mined from 7,793 annotated genes (as described in Figure 5C) and plotted as a histogram. The black arrow marks the 3’ UTR of *ck-1a* (NCU00685) at 1,739 bp in length (**A**). UTR lengths from *Drosophila melanogaster* were mined from 13,552 uniquely annotated genes (Ensembl GTF version BDGP6, accessed on 8/5/2020 from Illumina iGenomes) and plotted as a histogram. The black arrow marks the UTR of *dbt* (FBgn0002413) at 2,443 bp in length (**B**). UTR lengths from *Mus musculus* were mined from 20,477 uniquely annotated genes (Ensembl GTF version GRCm38, accessed on 8/5/2020 from Illumina iGenomes) and plotted as a histogram. The black arrow marks the UTR of CSNK1 D (ENSMUSG00000025162) at 2,157 bp in length, and the blue arrow corresponds to CSNK1 E (ENSMUSG00000022433) at 1,456 bp (**C**). UTR lengths from *Homo sapiens* were mined from 22,401 uniquely annotated genes (Ensembl GTF version GRCh37, accessed on 8/5/2020 from Illumina iGenomes) and plotted as a histogram. The black arrow marks the UTR of CSNK1 D (ENSG00000141551) at 2,113 bp in length, and the blue arrow corresponds to CSNK1E (ENSG00000213923) at 1,247 bp (**D**).

**Supplementary Table 1.**
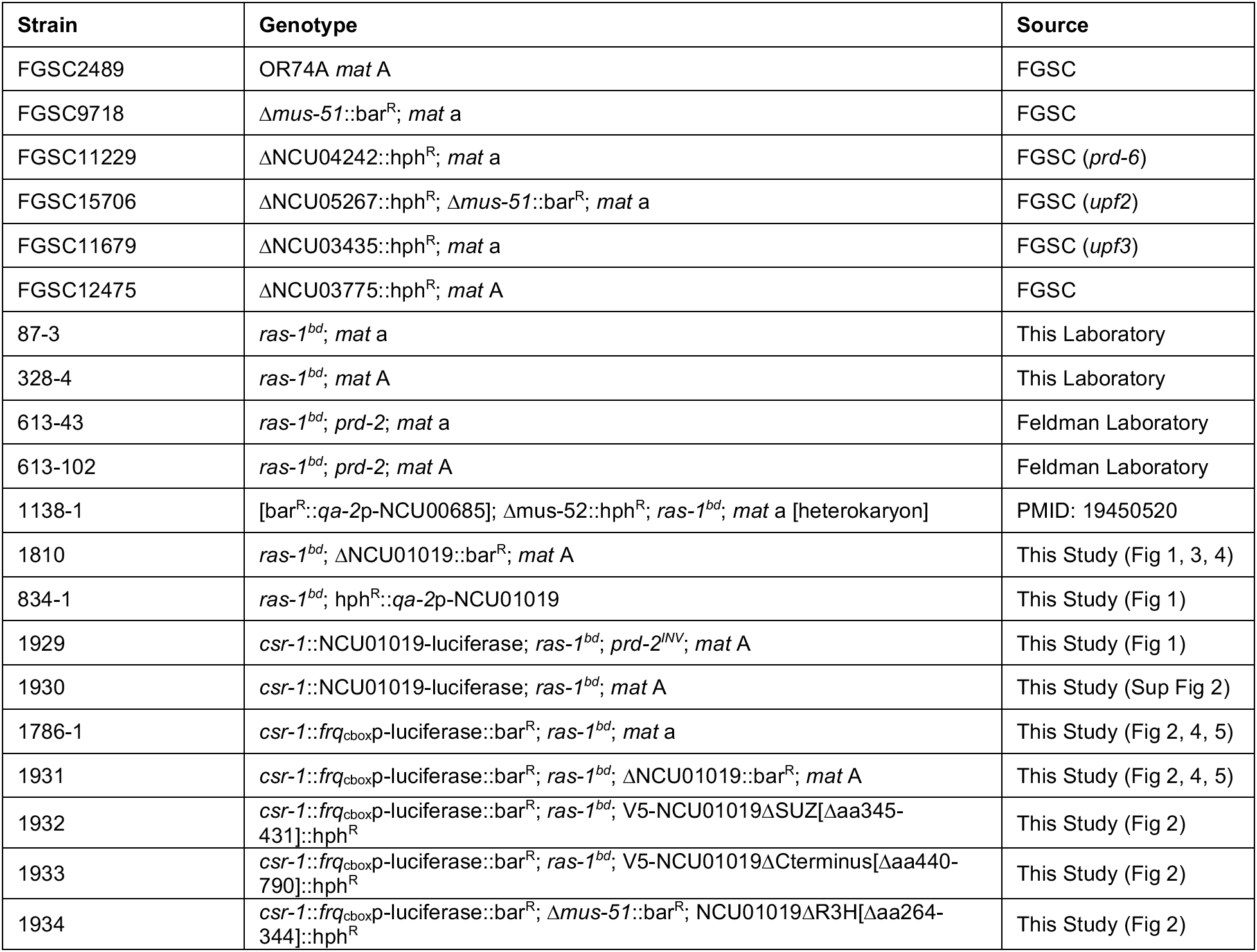
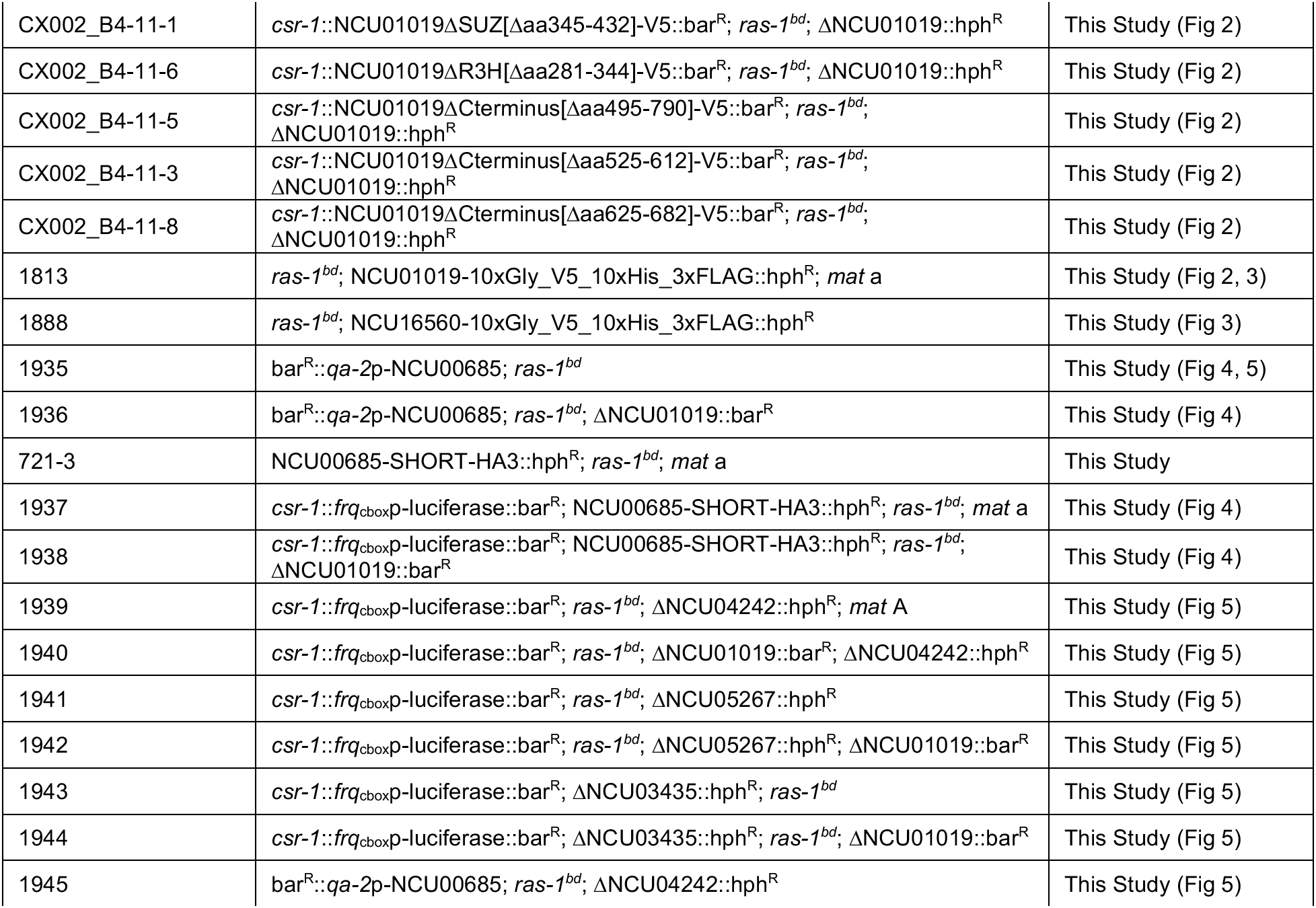
*Neurospora crassa* strains used in this study.

## Acknowledgements

We thank the Fungal Genetics Stock Center (Kansas City, Missouri, USA) for curating *N. crassa* strains. We thank Arun Mehra for discussions on preliminary work to identify the clock-relevant mechanism of the *prd-6* mutation, Bin Wang for assistance in constructing and validating the CKI^SHORT^ hyperactive allele, Jill Emerson for assistance in constructing *Δprd-2* (NCU01019), and Brad Bartholomai for discussions on *prd-2.* The *Neurospora* CK1a antibody was courtesy of Michael Brunner (University of Heidelberg).

